# Coevolution of longevity and female germline maintenance

**DOI:** 10.1101/2023.12.03.569746

**Authors:** Julian Baur, Mareike Koppik, Uros Savkovic, Mirko Dordevic, Biljana Stojkovic, David Berger

## Abstract

An often-overlooked aspect of life-history optimization is the allocation of resources to protect the germline and secure safe transmission of genetic information. While failure to do so renders significant fitness consequences in future generations, germline maintenance comes with substantial costs. Thus, germline allocation should trade-off with other life history decisions and be optimized in accordance with an organism’s reproductive schedule. Here we tested this hypothesis by studying germline maintenance in lines of seed beetle, selected for early (E) or late (L) reproduction for 350 and 240 generations, respectively. Female animals provide maintenance and screening of male gametes in their reproductive tract and oocytes. Here, we revealed the ability of young and aged E and L-females to provide this form of germline maintenance by mating them to males with ejaculates with artificially elevated levels of protein and DNA damage. We find that germline maintenance in E-females peaks at young age and then declines, while the opposite is true for L-females, in accordance with the age of reproduction in respective regime. These findings identify the central role of allocation to secure germline integrity in life history evolution and highlight how females can play a crucial role in mitigating effects of male germline decisions on mutation rate and offspring quality.

## Introduction

The interplay between growth, reproduction, and survival is a central focus of life history theory (1,2). Trade-offs among these three fundamental aspects of an organism’s life play key roles in dictating the evolution of reproductive schedules, with cascading effects on organismal aging and demography (3–5). Indeed, some species invest heavily in early reproduction at the expense of longevity, while others prioritize growth and survival before reproducing, and these alternative strategies are expected outcomes of differences in mortality risk and age-related changes in the strength of natural selection (1,2,6).

Another, often less explicitly defined, dimension of life history optimization involves the allocation of resources to protect the germline (7–9). The integrity of the germline is essential for the transmission of genetic information to subsequent generations, and hence, the evolutionary consequences of germline maintenance strategies can be considerable both at the individual and population level (10–12). The costs of germline maintenance may be substantial (8), in part due to the expensive repair machinery required to keep the rate of deleterious germline mutations at levels several magnitudes lower than the somatic mutation rate (13). This sets the stage for trade-offs between the germline and other life history traits that are governed by related processes, such as somatic maintenance and repair affecting longevity (14,15), or germline replication rate affecting sperm competition success (16–18). Thus, costly maintenance of the germline may secure the genetic quality of future generations at the cost of reduced reproduction and/or survival of the parent (7–9).

Theories of aging therefore do not only predict age-specific optimization of classic life history traits, but also concerted evolution of processes aimed at protecting the germline. For example, the mutation accumulation theory of aging (19), from hereon: “MA”, predicts a decline in germline maintenance past an organism’s reproductive peak due to weakened selection and the accumulation of alleles with deleterious effects late in life. Indeed, ageing not only leads to somatic deterioration (20), but also negatively affects the maintenance of the germline (9), shown, for example, by the fact that *de novo* germline mutation rates in primates seem to be foremost associated with advancing age (21–23). Ageing can also be driven by antagonistic pleiotropy (24), from hereon: “AP”, whereby alleles with beneficial effects on germline maintenance at the peak of reproduction (e.g. via increased allocation of energy to germline repair) are favoured by selection despite potential deleterious side-effects on somatic maintenance (15), reproduction (14,16), and/or germline maintenance at other ages.

There is now growing evidence that germline decisions can be studied through a life-history lens. For instance, germline maintenance strategies can differ between environments that place different needs on maintenance versus reproduction (25–31), or between individuals that differ in their overall energy budgets and allocation decisions (14,16,18,32,33).

Interestingly, female animals possess the ability to provide maintenance not only of their own germline cells, but also of male ejaculates and haploid male DNA, by synthesizing anti- oxidants in the reproductive tract to neutralize mutagenic reactive oxygen species, and by including mRNA and proteins in the oocyte to detect and repair DNA damage in the early zygote (34–36). Additionally, females may also exert cryptic female choice of male sperm that compete for fertilization of eggs inside the female reproductive tract (37,38), which can serve as an additional barrier screening against male gametes of low genetic quality, even among sperm within an ejaculate from a single male (39,40). This process bares similarity to apoptosis (controlled cell death) of damaged germline cells, which limits the passing on of deleterious mutations to offspring (36,41–44). Here we consider all these related processes (cryptic female choice, apoptosis, antioxidant defence, and DNA repair) as alternative and non-mutually exclusive ways that females can exert germline maintenance in the broad sense. This female control over male gamete quality sets the stage for an intricate evolutionary dynamic, where care over transferred ejaculates and newly formed zygotes is the ultimate decision of the female, but the need for it might be dictated by the germline decisions of the male partner, or, like in human populations, his age (45–47).

Even though there now is a burgeoning understanding of the mechanisms that cause age-dependent variation in DNA-repair and maintenance of the germline, little is known about the evolutionary potential of such mechanisms. In humans, for example, changes to environmental (48), pharmaceutical (49) and genetic (50) factors can increase longevity substantially within one or a few generations, and the peak of reproduction has in many populations changed drastically over the last century (51,52). However, the long-term genetic consequences of lifespan extension depend on whether selection on reproductive schedules are coupled with concomitant changes in germline repair and maintenance, which remain unexplored. Here we harnessed the power of long-term experimental evolution to investigate how the evolution of lifespan affects age-dependent germline maintenance in the seed beetle, *Acanthoscelides obtectus.* Using a set of lines that were established in 1989 and selected for early or late reproduction for more than 350 and 240 generations, respectively, we examined germline maintenance in young and aged females receiving male ejaculates with artificially inflated levels of free radicals and DNA and protein damage (Fig. 1). This allowed us to explore i) whether female germline maintenance evolves in concerted fashion with reproductive schedules and other life history traits, and ii) whether the AP (24) or the MA (19) theory of ageing best explains the evolutionary patterns observed.

**Figure 1:**
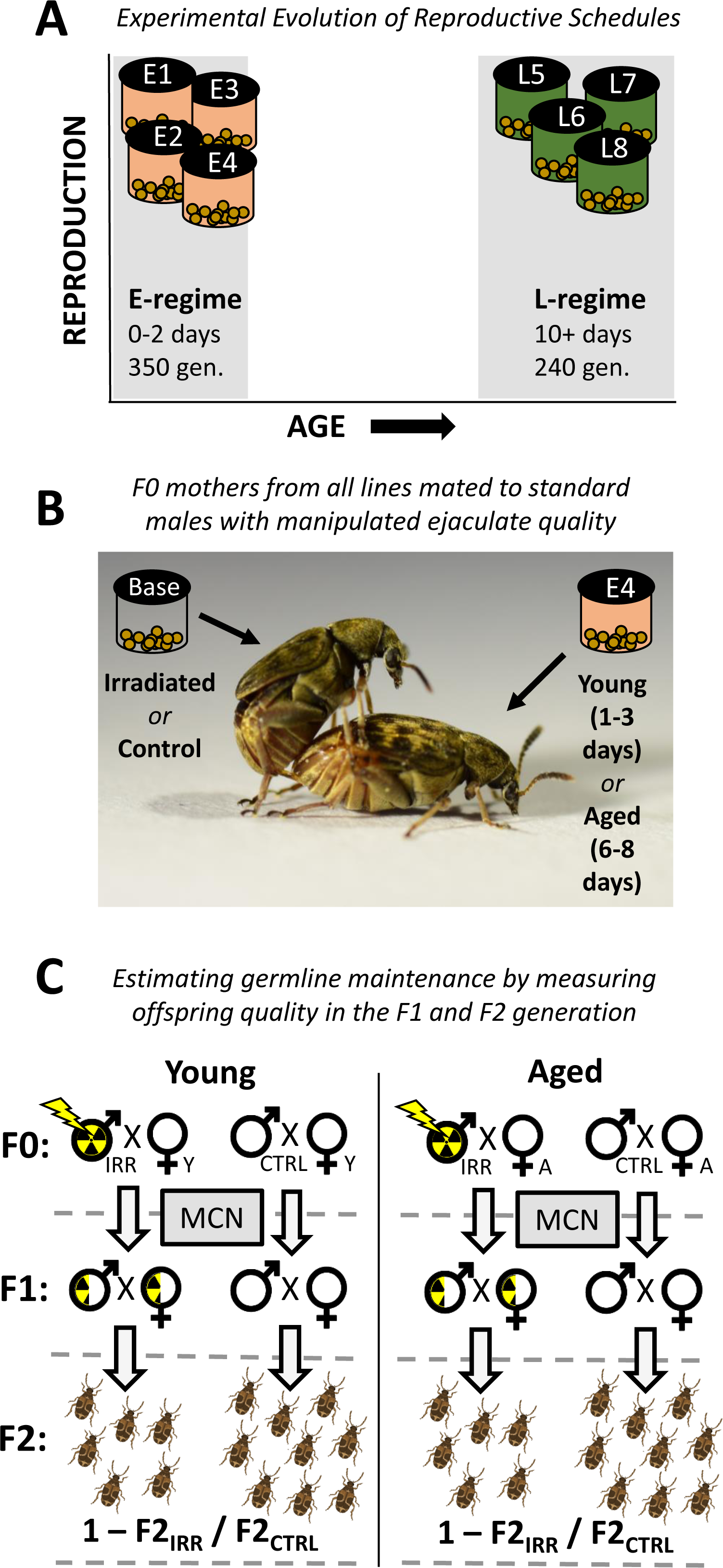
Experimental design. **A)** The experiment included eight experimental evolution lines; four from the E- (early reproduction; salmon) and four from the L- (late reproduction; green) regime. The E-regime was propagated by harvesting eggs laid after only 2 days of adult reproduction, whereas the L-regime was propagated by providing individuals that had aged 10 days with host seeds and harvesting the eggs. The E and L regime had been kept for 350 and 240 generations, respectively, prior to the experiment. **B)** Females from all lines, that were either young (1-3 days) or aged (6-8 days), were mated to standard males from the base population and allowed to lay eggs for 48h. The males were either serving as controls or they had been irradiated to generate a damaged ejaculate that would challenge maintenance in the female reproductive tract and oocytes. Offspring resulting from the matings to control males were counted to estimate age-dependent reproductive schedules of the F0 females in the experiment. **C)** F1 offspring were crossed within line and treatment group (while avoiding inbreeding) using a middle-class neighbourhood (MCN) design to relax selection against induced (epi)mutations. Counts of emerging F2 adult offspring were used to assess age-specific female germline maintenance by comparing the proportional reduction in offspring quality in lineages deriving from either young and aged F0 females, mated with either irradiated or control males. Photo credit for panel B: Mareike Koppik. Beetles in panel C were generated using BioRender.

## Methods

### Study species

The seed beetle, *Acanthoscelides obtectus* (Coleoptera; Chrysomelidae; Bruchidae) is a pest of stored legumes, foremost, the common bean, *Phaseolus vulgaris* (53). Females lay eggs either directly under or in close proximity of host seeds and the hatched larvae burrow into a seed where juvenile development takes place (54). After pupation, adults emerge from the seed and typically become reproductively active within 24 hours. Adult beetles are aphageous, meaning they dońt need nutrition or water to be able to reproduce and finish their lifecycle (53,55).

### Experimental populations

The original base population, from which the experimental evolution lines were derived, was created by mass-mating wild-caught beetles from three locations in the area of Belgrade, Serbia, in 1983 (54). Subsequently, the population was maintained on *P. vulgaris* in the laboratory for 3 years (27 generations) at large population size (ca. 5000 individuals), until the experimental evolution lines were created in 1986. Experimental evolution was performed in dark incubators at 30°C. No food or water was offered to adult beetles.

During experimental evolution, one line was maintained under the same conditions as the base population (from here on referred to as the “base” line); after 40 days of development, 400-500 adult beetles were chosen at random and provided with fresh host seeds and allowed to reproduce for the next generation. Each of four replicate lines selected for early reproduction (E-lines) was created by taking 400 adults at random on the peak of adult emergence. To propagate the next generation, these adults were transferred to a new bottle containing host seeds *ad libitum* and were there allowed to reproduce and lay eggs for 48h, after which all adults were removed. Another four replicate lines were selected for late reproduction (L-lines) by taking 100 newly emerged (0-24h) adults every day for up to 10 days. Each of these subgroups was kept in a vial without seeds for 10 days. On the 10^th^ day of each subgroup all the surviving beetles of a given replicate line were added to a single bottle containing fresh host seeds to propagate the next generation (Fig. 1A).

Development time of E-lines has evolved to be 29 days and the average virgin lifespan around 13 days in aphagous conditions, while L-lines emerge after 32 days and virgin beetles live on average 26 days for females and 22 days for males (56). The regimes have also diverged in other traits tied to rate-dependent life-history, such as the day of first oviposition (E: 1 day after adult eclosion, L: 4 days after adult eclosion) (57), metabolic rate, and body size (58). Care was taken not to select directly on development time (by collecting adult beetles around the peak of emergence in each generation) or body size (by providing host seeds ad libitum. It can thus be expected that life history evolution in the lines is a result of correlated responses to the applied selection on reproductive schedules. For more information on the maintenance of the experimental evolution lines, see (54,55).

### Experimental design

The experiment was started in 2020, after 240 (L-lines) and 350 (E-lines) generations of experimental evolution. To estimate the evolution of age-dependent germline maintenance we compared females of all lines mated to males when aged 1-3 days (young treatment) or 6-8 days (aged treatment) (Fig. 1B). Because we wanted to make inferences about evolved differences in biological aging and correlated life history between the regimes, we compared E and L females at the same chronological age. We did not pick an older age than 8 days since E females age rapidly and we wanted to avoid any bias in form of selective disappearance (59) of low quality individuals in the E-lines. We note, however, that 6-8 days is not truly an old age for L-lines, but rather represents a relatively young reproductive age (54,56). The comparison between E and L females of the same chronological age should thus be interpreted in relative terms.

All lines were reared in staggered fashion by setting up several rearing containers of each line in the two generations prior to the F0 generation of the experiment, so that emergence of females would happen continuously during the experiment and females assigned to the two age treatments could be assayed simultaneously and mated to standard males from the same cohort (see below). We collected females on the day of their emergence, randomly assigned them to the young or aged treatment, and then isolated them in perforated 0.5ml Eppendorf tubes and left them age. Females of both age treatments were then mated to young (1-2 days) virgin males from the base population in 60 mm petri-dishes placed on heating plates kept at 30°C. Couples were observed to avoid multiple mating. Males were removed after a mating behaviour had been observed and females were allowed to lay eggs in a new 60 mm petri-dish containing host seeds *ad libitum* for 48h after the mating.

The base males were either untreated (controls) or had been exposed to a 10 Gy dose of γ- radiation, which causes substantial cellular damage to the male germline (43) and impairs male fertility in seed beetles (14,16,18). The dose of 10 Gy was chosen based on our previous studies using similar experimental designs in the related seed beetle *Callosobruchus maculatus* (16,18,60), as well as another study on *A. obtectus* (61). Hence, the female repair, maintenance, and potential screening of damaged male gametes in each line could be assessed by comparing the offspring production in lineages stemming from females mated to control and irradiated males. γ-radiation produces free radicals with mutagenic properties and causes direct damage to DNA within sperm cells (43) and to proteins within the ejaculate (62). Hence, our protocol captures a multitude of effects that occur naturally in male ejaculates (8,63), but that will have been greatly amplified by the irradiation treatment.

As our interest lies in capturing effects of variation in female germline maintenance that transcend generations, we scored the fitness consequences of variation in female maintenance by counting the number of emerging F2 adults produced by all descending lineages. To achieve this, F1 males and females stemming from the same treatment group and line were paired (while avoiding sib-mating) and allowed to mate and produce F2 offspring in petri-dishes with ad libitum host seeds (Fig. 1C). The emerging F2 adult offspring were counted to estimate each couple’s fecundity, which was used to estimate germline maintenance of their F0 grandmothers (see below: “Statistical analysis”).

About half of the F0 couples produced no offspring (Supplementary Table 1a). This was likely due to two reasons. First, the mating latency of *A. obtectus* can be relatively long, and because the time in between male irradiation and mating needed to be standardized (maximum of 30 minutes), many couples did not mate in this time. Secondly, even when a mating was thought to have been observed, several couples did still not produce any offspring. Using a binomial model, we tested whether the incidence of zero fertility was related to any of the factors of interest in our experiment (i.e., irradiation treatment, selection regime, and age treatment). However, despite large sample size, no factor had a significant effect on infertility. Indeed, all effects including male irradiation treatment were very weak and non-significant (all P > 0.3), and the proportion of infertile couples was relatively equal across treatments (E: young ctrl = 0.43; young irr = 0.48; old ctrl = 0.59; old irr = 0.61. L: young ctrl = 0.50; young irr = 0.39; old ctrl = 0.48; old irr = 0.50). Hence, we carried out our downstream analyses based on the remaining data from the 830 F0 couples that produced offspring. Each line and age-class was represented by 10-20 females mated to control males, and 20-30 females mated to irradiated males. More couples were set up for the irradiated treatment group since the induced damage to male ejaculates was expected to generate more variance in offspring quality among lineages, requiring a larger sample size to result in the same precision of estimates as for the control lineages.

To reduce the effect of selection against epigenetic and DNA mutations induced by the irradiation we applied a Middle-Class Neighbourhood (MCN) design (64) (Fig. 1C). The MCN design restricts each F0 couple to contribute the same number of F1 offspring for future study, irrespective of their overall productivity. We note, however, that F0 couples that produced less than four F1 offspring were not represented in the final data, and hence, selection on very strongly deleterious effects of irradiation could not be excluded by the MCN approach. For those F0 couples that did produce offspring, we aimed at propagating a single male and female from each couple, so that each F0 couple would replace itself.

However, due to variability in the timing of eclosion of F1 individuals, additional F0 couples fell out of the experiment as we did not want to age F1 beetles before setting them up in assays. These strict criteria resulted in 490 F1 couples being analysed for offspring production. Each line and age class was represented by ca 11-27 and 5-15 F1 couples that originated from irradiated and control lineages, respectively (Supplementary Table 1b).

The experiment was conducted in three temporal blocks, 3 days apart, corresponding to the three occasions on which we irradiated F0 males from the base population (Supplementary Table 1a). In the F1 generation, when offspring productivity assays were set up, there was considerable overlap between these blocks due to variable timing of egg laying and juvenile development time, so that combinations of F1 individuals stemming from all combinations of F0 dates were crossed. Hence, we did not include the effect of irradiation day in further analysis.

### Statistical analysis

We analyzed data using generalized linear mixed effect models implemented in the package MCMCglmm (65) for R (66). Offspring counts were analysed assuming Poisson distributed error. We used flat and uninformative priors for all models. We ran all models for 1.05 million iterations with an initial burn in of 50k iterations and a thinning factor of 500 to avoid autocorrelations, resulting in a posterior sample of 2000. P-values for all comparisons were calculated based on model posterior distributions, where significance of main effects was calculated by comparing posteriors of marginal means of two given groups (e.g. overall difference in offspring production in irradiated and control lineages). We used ggplot2 for graphical illustration (67).

To provide an estimate of the extent of age-specific optimization of reproductive schedules at the current stage of experimental evolution, we first compared counts of offspring produced by young and aged F0 females of all E- and L-lines over the 48h period, when mated to the standard base males belonging to the control (un-irradiated) treatment (N = 307). This model included female age treamtent (young vs. aged), evolution regime (E vs. L), and their interaction, as fixed effects of interest. We also added the main effect of male age (1 or 2 days old) as a blocking factor. Replicate line, crossed with age, were added as random effects (model summary in Supplementary Table 2a).

To analyse differences in female germline maintenance we compared the F2 offspring counts of control and irradiated lineages. This model included fully crossed fixed effects of evolution regime (E vs. L), F0 female age treatment (young vs. aged) and male radiation treatment (control vs. irradiated). Note that the three-way interaction between these effects tests our focal hypothesis that the E- and L-regime have diverged in age-specific female germline maintenance. Replicate line crossed with age treatment and radiation treatment were included as random effects (model summary in Supplementary Table 2b). To provide an intuitive measure of germline maintenance for reporting effect sizes, we also calculated the relative reduction in fertility due to unattended damage in the male ejaculate for each replicate line for young and aged females, according to: Δζ = 1 - ζ_IRR, *i*_ / ζ_CTRL, *i*_, where ζ_IRR, *i*_ and ζ_CTRL, *i*_ is the number of F2 offspring produced by lineages deriving from F0 females of age treatment *i*, mated to irradiated and control males, respectively (see: Fig. 3B).

## Results

### Evolution of age-specific reproductive schedules

There was no difference in F1 offspring production between young E and L females mated to control males in the F0 generation (p_MCMC_ = 0.28). However, there were clear differences in aging. As expected, aged females originating from the E-regime showed a sharp decline in offspring production (p_MCMC_ < 0.001), while no such effect was seen in the aged L-females. As follows, the interaction between regime and female age treatment was highly significant (p_MCMC_ < 0.001) and accorded with predictions (Fig. 2).

**Figure 2:**
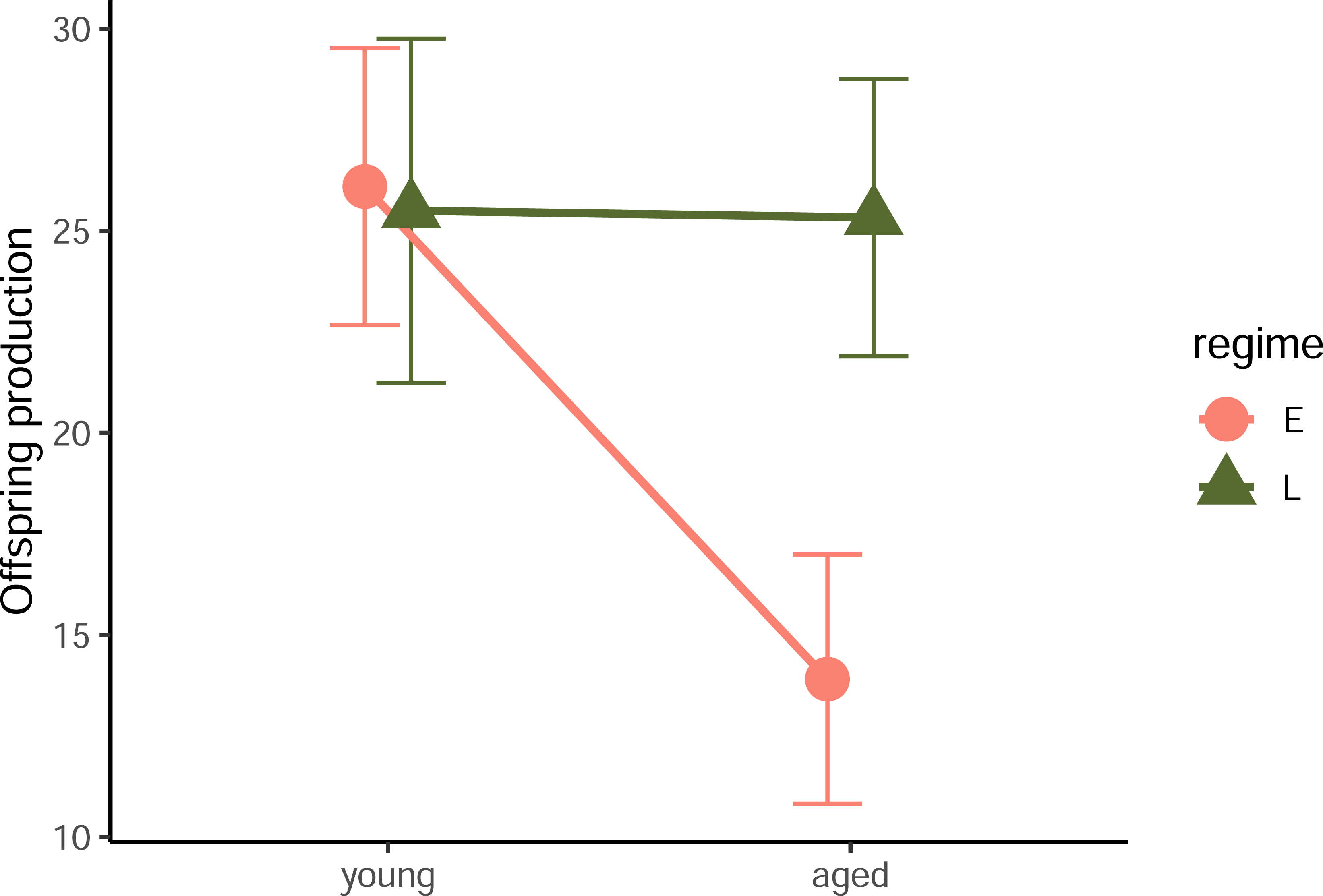
Evolution of female reproductive schedules. The number of F1 offspring produced by young and aged F0 females of the E (salmon) and L (green) regime, during the 48h of egg laying following mating to standard control males from the base line. Shown are means and 95% confidence limits.

**Figure 3:**
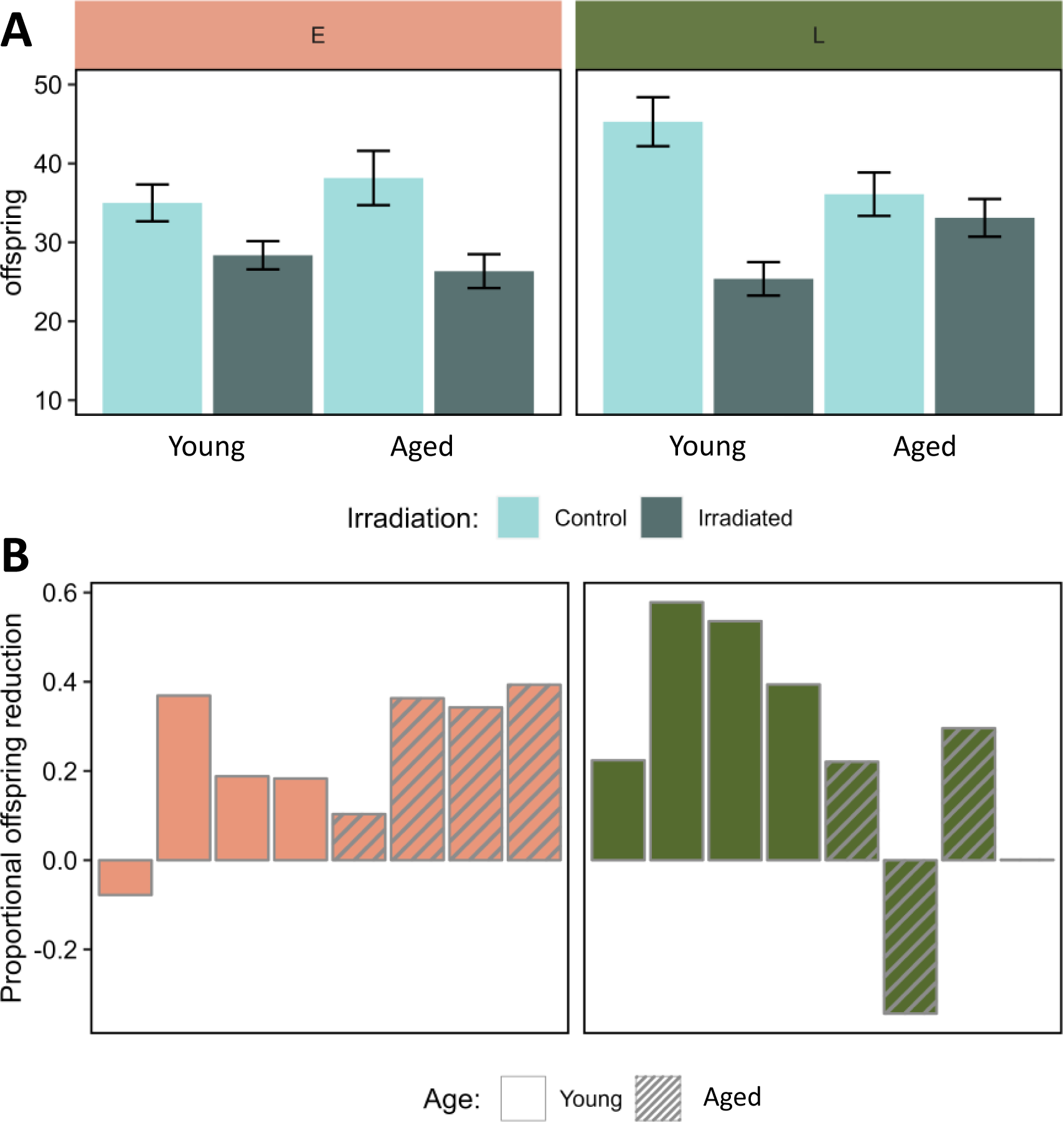
Evolution of age-dependent female germline maintenance. **A)** F2 offspring production in lineages stemming from F0 grandmothers mated to either a control (light bars) or irradiated (dark bars) male. Grandmothers were either young (1-3 days) or aged (6-8 days) at mating, and belonged to either the early reproducing (E: salmon, left panel) or late reproducing (L: green, right panel) evolution regime. Shown are means and standard errors. **B)** Line-specific estimates of the relative reduction in F2 offspring production induced by the irradiation treatment in F0 males from the base population, calculated as : Δζ = 1 - ζ_IRR, *i*_ / ζ_CTRL, *i*_, where ζ_IRR, *i*_ and ζ_CTRL, *i*_ is the number of F2 offspring produced by lineages deriving from F0 females of age *i*, mated to irradiated and control males, respectively. This estimate is thus inversely related to the efficiency of germline maintenance in F0 mothers. Plain and striped bars represent lineages originating from young and aged F0 grandmothers, respectively.

### Coevolution of reproductive schedules and female germline maintenance

The irradiation treatment led to an overall decrease in F2 offspring numbers (effect of irradiation in L-regime: p_MCMC_ < 0.001; effect of irradiation in E-regime: p_MCMC_ = 0.002). Taken across both age treatments, there was no difference between evolution regimes in the effect of irradiation on offspring production (regime*irradiation interaction: p_MCMC_ = 0.74), providing no evidence for strong divergence in female germline maintenance overall. However, in terms of the age-dependency, the two regimes seemed to behave very differently (regime*age*irradiation interaction; p_MCMC_ = 0.008, Fig. 3). This difference was due to that the effect of irradiation was stronger in young, compared to aged, L-females (p_MCMC_ = 0.004), while the opposite pattern was observed for E-females (although the effect of age was not significant, p_MCMC_ = 0.35). These results confirm that female germline maintenance has evolved age-specific optimization.

## Discussion

Here we have shown that selection on reproductive schedules can lead to the evolution of age-specific female germline maintenance that mitigates deleterious effects caused by low quality male gametes. Such germline maintenance is a somewhat underestimated form of parental care with genetic consequences that can transcend generations. Our findings echo previous studies that have provided evidence that germline maintenance is costly and can trade off with life-history traits (14–16,32,68), emphasizing that germline maintenance should be considered a life-history trait in its own right (7,8,14,69–71).

*Did germline maintenance evolve via mutation accumulation or antagonistic pleiotropy?* The observed evolution of aging follows the general expectation that repair and maintenance in the female reproductive tract evolves to optimize function at peak reproduction. The mutation accumulation (MA) theory of ageing (19) predicts that E-lines should show efficient maintenance early in life, but a sharp decline in maintenance at old age, as old E-beetles have not been exposed to selection for >350 generations. In L-lines, efficient maintenance of both young and aged females is expected, given that mature gametes have already formed at adult eclosion (61) and should need continuous maintenance. Indeed, the observed reproductive schedules of young and aged E- and L- females mated to control (unirradiated) males were qualitatively consistent with the MA hypothesis (Fig. 2). However, for female germline maintenance, E-females showed only a modest reduction with increasing age, and instead it was young L-females that showed the lowest maintenance (Fig. 3). L- females have not been provided with egg-laying substrate at young age for over 240 generations and egg-laying at this age thus represents a novel condition, which means that accumulation of mutations with conditional deleterious effects only at young age cannot be excluded. However, while the dip in germline maintenance was larger in young (relative to aged) L-females compared to the dip in aged (relative to young) E-females, the E-regime had undergone more generations of (potential) accumulation of mutations with age-specific effect. Thus, the observed patterns of aging of germline maintenance are not fully compatible with the MA hypothesis.

Another (not mutually exclusive) explanation for the evolved aging trajectories is antagonistic pleiotropy (AP) (24). Indeed, previous findings have shown that age-related changes in fertility and longevity in the E- and L lines of *A. obtectus* seem to be a combined result of both MA and AP (72). Pleiotropic effects of genes improving germline maintenance at young age could lead to reproductive disadvantages at later ages (8,73) and/or reduced survival and longevity via trade-offs between somatic and germline DNA-maintenance (8,15), and hence, would be disfavoured by selection in the L-regime. Interestingly, the pronounced dip in germline maintenance in young L-females is also mirrored by common observations of reduced fertility due to reproduction too early in life across animal taxa (74). Here it is also worth underlining that the aged treatment in L-females actually represents a young reproductive age in this evolution regime, that likely corresponds to the onset of female reproductive investment during experimental evolution (54,55). Hence, the E- females in the young treatment and L-females of the aged treatment should closely correspond to each other in terms of their biological age, and indeed, also show similar levels of reproductive output (Fig. 2) and germline maintenance (Fig. 3).

### Male-female coevolution

The potential for females to care over male ejaculates already at the very onset of fertilization opens up interesting eco-evolutionary scenarios and the potential for sexual conflict over parental care, even in species traditionally assigned as having little to no care (18,68,75,76). For example, in most species with internal fertilization, healthy females may efficiently compensate for poor ejaculates of older males, while the converse seems unlikely. This asymmetry should have implications for assortative mating by age. For example, the baseline germline mutation rate is several times greater in males compared to females in most animal taxa (71,77,78) and, at least in well-studied groups like primates, this bias increases with age (79,80). Thus, mating decisions based on partner age are likely to have considerable consequences, and the form of maternal care studied here should be intertwined in those decisions.

Germline maintenance decisions can also depend on the overall condition of the individual and state-dependent allocation decisions between competing life-history demands (8,9,14,33), with implications for sexual selection theory and mate choice processes (16,18,81,82). Viewing germline maintenance through a resource acquisition and allocation lens (83) suggests that optimal mate choice is contingent on not only choosers’ inferences of genetic quality in offspring based on the phenotype of their mating partner, but also of how this phenotype influences changes to the genetic quality of its gametes. For example, in many polyandrous species, strong postcopulatory sexual selection (sperm competition) can favour compromised germline maintenance in males (16–18,69). This suggests that males that are most successful in reproductive competition may sometimes pass on the greatest load of mutations to their offspring, questioning why females should prefer them in the first place (16,81,82)? Interestingly, however, optimality theory predicts that females my tailor care depending on the assessed state of male gametes (76), with the result that a male’s mutation rate is not independent of his female mate and her potential compensation for his short-comings, which can complicate interpretations of both age- and sex-biases in mutation rate (45).

Another way that females could influence the resulting quality of their offspring is via haploid selection through cryptic female choice of male sperm (40). In this study, we defined female germline maintenance broadly, including both care of male gametes in form of DNA repair and antioxidant defence, as well as screening against damaged sperm via cryptic female choice or damaged zygotes via apoptosis. Our study does not allow us to discriminate among these hypotheses, that are not mutually exclusive (42–44). Yet, it is interesting to note that, while all these processes equate to germline maintenance and increased offspring quality from a female’s perspective, only female care improves male gamete quality, while female screening eliminates the male’s gametes. Therefore, the type of female maintenance may have different implications for male fitness, with the level of sperm competition within and between ejaculates of different males a predicted driver of these dynamics (84,85). We do not know the extent of sperm competition in the E- and L- regime, although it would seem likely that the number of different male mates per female might be higher in L-females due to their prolonged lifespan. Nevertheless, we found no obvious differences between the evolution regimes in germline maintenance overall, or at the peak of early reproduction (young E females versus aged L-females) (Fig. 3). Future research could investigate if females kept under high levels of sperm competition evolve increased cryptic female choice and if this is associated with correlated evolutionary responses in other aspects of germline maintenance.

In relation to male-female coevolution, we note that there is a possibility that our results could have been driven by an indirect effect of female reproductive aging. If the F0 reference males in our experiment could assess the reproductive status of females, it is possible that males may have preferred females at their reproductive peak, and allocated resources in their ejaculates accordingly, resulting in higher offspring quality in young E- and aged L-females. However, we find this hypothesis unlikely due to three reasons. First, in addition to requiring the ability of males to assess variation in female condition, there would also need to be sufficient fitness gains for this male allocation decision to evolve. We find this unlikely given the high-density scramble mating system under which the reference males have been evolving for at least 300 generations in the lab environment, and the relatively minor effects on female fertility caused by large variation in male ejaculate volume and quality across other species of seed beetle (86). Second, the allocation decision would have needed to be very fast as most F0 males mated within less than X minutes after have being introduced to the females, while processes that affect DNA repair and oxidant defence of male sperm typically occur over longer time frames (43), as does sperm maturation in the relative *C. maculatus* (16). Third, this effect would also need to take effect in irradiated and not control lineages to generate the observed pattern (Fig. 3).

Nevertheless, our current data do not allow us to completely rule out that this effect could have contributed to our results.

## Conclusions

Can mechanisms that protect the germline evolve fast enough to keep up with rapid environmental and demographic change? This question is of importance for understanding long-term evolutionary responses that depend on both demographic parameters and the germline mutation rate. It is also a question tied to ongoing debates in human reproductive biology against the background of the increasing age at first reproduction in western civilisations (87,88). While assisted reproduction technologies may help to mitigate immediate age-related fertility declines associated with such changes, less attention is allotted to the evolutionary implications of ever later reproduction in human populations. Our study provides compelling evidence that female care and/or screening of male gametes can evolve in response to selection on reproductive schedules. Germline maintenance peaked at the time of reproduction in E- and L-lines, and the observed evolution of aging of the germline is compatible with some role for antagonistic pleiotropy in the underlying genes. This suggest that balancing selection on concerted life-history syndromes might be responsible for the maintenance of genetic variation in germline maintenance in the wild founding population, which likely contributed to the observed evolutionary response. Our study highlights the evolutionary importance of this, often underappreciated, type of maternal care and we hope to stimulate further research aimed at elucidating the processes that govern the coevolution of germline maintenance, mate choice processes, and other life-history traits.

## Data accessibility

All data and code accompanying this study is uploaded to Dryad […].

## Author contributions

DB, JB and MK designed the study. JB and MK performed the study. JB and DB analyzed the data, and DB and JB wrote the paper. US, MD and BS maintained, provided and prepared the experimental evolution lines. All authors read and commented on drafts of the manuscript.

## Acknowledgements

We thank Johanna Liljestrand-Rönn for help in the lab, the kind people at BMC Uppsala for providing access to the radiation source, and the members of the seed beetle lab group for helpful discussions. The maintenance of the experimental *A. obtectus* populations was financed by grant from the Ministry of Science, Technological Development and Innovation of the Republic of Serbia, grant number 451-03-47/2023-01/200007. This study was supported by grant no. 2019-05023 from the Swedish Research Council (VR) to DB.

## References

1. Roff D. Evolution Of Life Histories: Theory and Analysis. Springer Science & Business Media; 1993. 554 p.

2. Stearns SC. THE EVOLUTION OF LIFE HISTORIES.

3. Flatt T, Heyland A, editors. Mechanisms of life history evolution: the genetics and physiology of life history traits and trade-offs. Oxford ; New York: Oxford University Press; 2011. 478 p. (Oxford biology).

4. Monaghan P, Charmantier A, Nussey DH, Ricklefs RE. The evolutionary ecology of senescence. Funct Ecol [Internet]. 2008 Jun [cited 2019 Jan 17];22(3):371–8. Available from: http://doi.wiley.com/10.1111/j.1365-2435.2008.01418.x

5. Promislow DEL, Tatar M. Mutation and senescence: where genetics and demography meet. Genetica [Internet]. 1998 Mar 1 [cited 2020 Oct 5];102(0):299–314. Available from: 10.1023/A:1017047212008

6. Lemaître JF, Moorad J, Gaillard JM, Maklakov AA, Nussey DH. A unified framework for evolutionary genetic and physiological theories of aging. PLOS Biol [Internet]. 2024 Feb 27 [cited 2024 Feb 29];22(2):e3002513. Available from: https://journals.plos.org/plosbiology/article?id=10.1371/journal.pbio.3002513

7. Avila P, Lehmann L. Life history and deleterious mutation rate coevolution. J Theor Biol [Internet]. 2023 Sep 21 [cited 2024 Feb 5];573:111598. Available from: https://www.sciencedirect.com/science/article/pii/S0022519323001959

8. Maklakov AA, Immler S. The Expensive Germline and the Evolution of Ageing. Curr Biol [Internet]. 2016 Jul [cited 2019 Jan 17];26(13):R577–86. Available from: https://linkinghub.elsevier.com/retrieve/pii/S0960982216303347

9. Monaghan P, Metcalfe NB. The deteriorating soma and the indispensable germline: gamete senescence and offspring fitness. Proc R Soc B Biol Sci. 2019 Dec 18;286(1917):20192187.

10. Agrawal AF, Whitlock MC. Mutation Load: The Fitness of Individuals in Populations Where Deleterious Alleles Are Abundant. Annu Rev Ecol Evol Syst [Internet]. 2012 Dec [cited 2018 Feb 19];43(1):115–35. Available from: http://www.annualreviews.org/doi/10.1146/annurev-ecolsys-110411-160257

11. Lynch M. Evolution of the mutation rate. Trends Genet [Internet]. 2010 Aug 1 [cited 2018 Aug 4];26(8):345–52. Available from: https://www.cell.com/trends/genetics/abstract/S0168-9525(10)00103-4

12. Muller HJ. Our load of mutations. Am J Hum Genet [Internet]. 1950 Jun [cited 2021 Jan 28];2(2):111–76. Available from: https://www.ncbi.nlm.nih.gov/pmc/articles/PMC1716299/

13. Lynch M. The Cellular, Developmental and Population-Genetic Determinants of Mutation-Rate Evolution. Genetics [Internet]. 2008 Oct [cited 2022 Jan 31];180(2):933–43. Available from: https://www.ncbi.nlm.nih.gov/pmc/articles/PMC2567392/

14. Berger D, Stångberg J, Grieshop K, Martinossi-Allibert I, Arnqvist G. Temperature effects on life- history trade-offs, germline maintenance and mutation rate under simulated climate warming. Proc R Soc B [Internet]. 2017 Nov 15 [cited 2018 Sep 19];284(1866):20171721. Available from: http://rspb.royalsocietypublishing.org/content/284/1866/20171721

15. Chen H yen, Jolly C, Bublys K, Marcu D, Immler S. Trade-off between somatic and germline repair in a vertebrate supports the expensive germ line hypothesis. Proc Natl Acad Sci. 2020 Apr 21;117(16):8973–9.

16. Baur J, Berger D. Experimental evidence for effects of sexual selection on condition-dependent mutation rates. Nat Ecol Evol [Internet]. 2020 May [cited 2020 Jul 10];4(5):737–44. Available from: https://www.nature.com/articles/s41559-020-1140-7

17. Blumenstiel JP. Sperm competition can drive a male-biased mutation rate. J Theor Biol [Internet]. 2007 Dec [cited 2019 Jan 17];249(3):624–32. Available from: https://linkinghub.elsevier.com/retrieve/pii/S0022519307004109

18. Koppik M, Baur J, Berger D. Increased male investment in sperm competition results in reduced maintenance of gametes. PLOS Biol [Internet]. 2023 Apr 4 [cited 2023 Apr 5];21(4):e3002049. Available from: https://journals.plos.org/plosbiology/article?id=10.1371/journal.pbio.3002049

19. Medawar PB. An unsolved problem of biology. Published for the College by H.K. Lewis; 1952.

20. Partridge L, Barton NH. On measuring the rate of ageing. Proc R Soc Lond B Biol Sci. 1996 Oct 22;263(1375):1365–71.

21. Kong A, Frigge ML, Masson G, Besenbacher S, Sulem P, Magnusson G, et al. Rate of de novo mutations and the importance of father’s age to disease risk. Nature. 2012 Aug;488(7412):471– 5.

22. Rahbari R, Wuster A, Lindsay SJ, Hardwick RJ, Alexandrov LB, Al Turki S, et al. Timing, rates and spectra of human germline mutation. Nat Genet. 2016 Feb;48(2):126–33.

23. Venn O, Turner I, Mathieson I, de Groot N, Bontrop R, McVean G. Strong male bias drives germline mutation in chimpanzees. Science [Internet]. 2014 Jun 13 [cited 2019 Jan 17];344(6189):1272–5. Available from: http://www.sciencemag.org/cgi/doi/10.1126/science.344.6189.1272

24. Williams GC. Pleiotropy, Natural Selection, and the Evolution of Senescence. Evolution [Internet]. 1957 [cited 2019 Mar 6];11(4):398–411. Available from: https://www.jstor.org/stable/2406060

25. 25. Berger D, Stångberg J, Baur J, Walters RJ. Elevated temperature increases genome-wide selection on de novo mutations. Proc R Soc B Biol Sci [Internet]. 2021 Feb 10 [cited 2021 Feb 3];288(1944):20203094. Available from: https://royalsocietypublishing.org/doi/full/10.1098/rspb.2020.3094

26. Chu XL, Zhang BW, Zhang QG, Zhu BR, Lin K, Zhang DY. Temperature responses of mutation rate and mutational spectrum in an Escherichia coli strain and the correlation with metabolic rate. BMC Evol Biol [Internet]. 2018 Aug 29 [cited 2018 Oct 29];18(1):126. Available from: 10.1186/s12862-018-1252-8

27. 27. Crean AJ, Immler S. Evolutionary consequences of environmental effects on gamete performance. Philos Trans R Soc B Biol Sci [Internet]. 2021 Jun 7 [cited 2021 Jun 21];376(1826):20200122. Available from: https://royalsocietypublishing.org/doi/abs/10.1098/rstb.2020.0122

28. Drake JW. Avoiding dangerous missense: thermophiles display especially low mutation rates. PLoS Genet. 2009;5(6):e1000520.

29. Kurhanewicz NA, Dinwiddie D, Bush ZD, Libuda DE. Elevated Temperatures Cause Transposon- Associated DNA Damage in C. elegans Spermatocytes. Curr Biol [Internet]. 2020 Dec 21 [cited 2023 Mar 3];30(24):5007-5017.e4. Available from: https://www.sciencedirect.com/science/article/pii/S0960982220314202

30. Puurtinen M, Elo M, Jalasvuori M, Kahilainen A, Ketola T, Kotiaho JS, et al. Temperature- dependent mutational robustness can explain faster molecular evolution at warm temperatures, affecting speciation rate and global patterns of species diversity. Ecography [Internet]. 2016 Nov [cited 2018 Jan 29];39(11):1025–33. Available from: http://doi.wiley.com/10.1111/ecog.01948

31. Svetec N, Cridland JM, Zhao L, Begun DJ. The Adaptive Significance of Natural Genetic Variation in the DNA Damage Response of Drosophila melanogaster. PLOS Genet [Internet]. 2016 Mar [cited 2018 Aug 4];12(3):e1005869. Available from: http://journals.plos.org/plosgenetics/article?id=10.1371/journal.pgen.1005869

32. Agrawal AF, Wang AD. Increased transmission of mutations by low-condition females: evidence for condition-dependent DNA repair. PLoS Biol. 2008 Feb;6(2):e30.

33. Sharp NP, Agrawal AF. Evidence for elevated mutation rates in low-quality genotypes. Proc Natl Acad Sci [Internet]. 2012 Apr 17 [cited 2019 Jan 17];109(16):6142–6. Available from: http://www.pnas.org/cgi/doi/10.1073/pnas.1118918109

34. Horta F, Catt S, Ramachandran P, Vollenhoven B, Temple-Smith P. Female ageing affects the DNA repair capacity of oocytes in IVF using a controlled model of sperm DNA damage in mice. Hum Reprod [Internet]. 2020 Mar 27 [cited 2021 Jul 7];35(3):529–44. Available from: 10.1093/humrep/dez308

35. Jaroudi S, Kakourou G, Cawood S, Doshi A, Ranieri DM, Serhal P, et al. Expression profiling of DNA repair genes in human oocytes and blastocysts using microarrays. Hum Reprod. 2009 Oct 1;24(10):2649–55.

36. Stringer JM, Winship A, Liew SH, Hutt K. The capacity of oocytes for DNA repair. Cell Mol Life Sci. 2018 Aug 1;75(15):2777–92.

37. Firman RC, Gasparini C, Manier MK, Pizzari T. Postmating Female Control: 20 Years of Cryptic Female Choice. Trends Ecol Evol [Internet]. 2017 May [cited 2021 Aug 11];32(5):368–82. Available from: https://www.ncbi.nlm.nih.gov/pmc/articles/PMC5511330/

38. Kustra MC, Alonzo SH. The coevolutionary dynamics of cryptic female choice. Evol Lett [Internet]. 2023 Aug 1 [cited 2023 Oct 13];7(4):191–202. Available from: 10.1093/evlett/qrad025

39. Alavioon G, Hotzy C, Nakhro K, Rudolf S, Scofield DG, Zajitschek S, et al. Haploid selection within a single ejaculate increases offspring fitness. Proc Natl Acad Sci [Internet]. 2017 Jul 25 [cited 2021 Feb 4];114(30):8053–8. Available from: https://www.pnas.org/content/114/30/8053

40. Immler S, Otto SP. The Evolutionary Consequences of Selection at the Haploid Gametic Stage. Am Nat [Internet]. 2018 Jun 8 [cited 2018 Sep 14];192(2):241–9. Available from: https://www.journals.uchicago.edu/doi/abs/10.1086/698483

41. Bhagavan NV, Ha CE. Chapter 22 - DNA Replication, Repair, and Mutagenesis. In: Bhagavan NV, Ha CE, editors. Essentials of Medical Biochemistry (Second Edition) [Internet]. San Diego: Academic Press; 2015 [cited 2020 Dec 15]. p. 401–17. Available from: http://www.sciencedirect.com/science/article/pii/B9780124166875000221

42. Borràs-Fresneda M, Barquinero JF, Gomolka M, Hornhardt S, Rössler U, Armengol G, et al. Differences in DNA Repair Capacity, Cell Death and Transcriptional Response after Irradiation between a Radiosensitive and a Radioresistant Cell Line. Sci Rep [Internet]. 2016 Jun 1 [cited 2019 Jun 4];6:27043. Available from: https://www.nature.com/articles/srep27043

43. Friedberg EC, Walker GC, Siede W, Wood RD. DNA Repair and Mutagenesis. American Society for Microbiology Press; 2005. 2845 p.

44. González-Marín C, Gosálvez J, Roy R. Types, Causes, Detection and Repair of DNA Fragmentation in Animal and Human Sperm Cells. Int J Mol Sci [Internet]. 2012 Oct 31 [cited 2019 Jan 17];13(12):14026–52. Available from: http://www.mdpi.com/1422-0067/13/11/14026

45. Gao Z, Moorjani P, Sasani TA, Pedersen BS, Quinlan AR, Jorde LB, et al. Overlooked roles of DNA damage and maternal age in generating human germline mutations. Proc Natl Acad Sci [Internet]. 2019 May 7 [cited 2019 Oct 2];116(19):9491–500. Available from: https://www.pnas.org/content/116/19/9491

46. Rahbari R, Wuster A, Lindsay SJ, Hardwick RJ, Alexandrov LB, Al Turki S, et al. Timing, rates and spectra of human germline mutation. Nat Genet [Internet]. 2016 Feb [cited 2022 Jan 17];48(2):126–33. Available from: https://www.nature.com/articles/ng.3469

47. Ségurel L, Wyman MJ, Przeworski M. Determinants of Mutation Rate Variation in the Human Germline. Annu Rev Genomics Hum Genet [Internet]. 2014 [cited 2019 Jan 18];15(1):47–70. Available from: 10.1146/annurev-genom-031714-125740

48. Klass MR. Aging in the nematode Caenorhabditis elegans: Major biological and environmental factors influencing life span. Mech Ageing Dev. 1977 Jan 1;6:413–29.

49. Johnson SC, Rabinovitch PS, Kaeberlein M. mTOR is a key modulator of ageing and age-related disease. Nature. 2013 Jan;493(7432):338–45.

50. Guarente L, Kenyon C. Genetic pathways that regulate ageing in model organisms. Nature. 2000 Nov;408(6809):255–62.

51. Khandwala YS, Zhang CA, Lu Y, Eisenberg ML. The age of fathers in the USA is rising: an analysis of 168 867 480 births from 1972 to 2015. Hum Reprod [Internet]. 2017 Oct 1 [cited 2022 Nov 11];32(10):2110–6. Available from: 10.1093/humrep/dex267

52. Mills M, Rindfuss RR, McDonald P, te Velde E, on behalf of the ESHRE Reproduction and Society Task Force. Why do people postpone parenthood? Reasons and social policy incentives. Hum Reprod Update [Internet]. 2011 Nov 1 [cited 2022 Nov 11];17(6):848–60. Available from: 10.1093/humupd/dmr026

53. Thakur DR. Taxonomy, Distribution and Pest Status of Indian Biotypes of Acanthoscelides obtectus (Coleoptera: Chrysomelidae: Bruchinae) - A New Record. Pakistan J Zool. 2012;44(1):189–95.

54. Tucić N, Gliksman I, Šešlija D, Milanović D, Mikuljanac S, Stojković O. Laboratory evolution of longevity in the bean weevil (Acanthoscelides obtectus). J Evol Biol. 1996;9(4):485–503.

55. Tucić N, Stojković O, Gliksman I, Milanović D, Šešlija D. Laboratory Evolution of Life-History Traits in the Bean Weevil (Acanthoscelides obtectus): The Effects of Density-Dependent and Age- Specific Selection. Evolution. 1997;51(6):1896–909.

56. Đorđević M, Savković U, Lazarević J, Tucić N, Stojković B. Intergenomic Interactions in Hybrids Between Short-Lived and Long-Lived Lines of a Seed Beetle: Analyses of Life History Traits. Evol Biol [Internet]. 2015 Dec 1 [cited 2023 Nov 13];42(4):461–72. Available from: 10.1007/s11692-015-9340-9

57. Tucić N, Šešlija D, Stanković V. The short-term and long-term effects of parental age in the bean weevil (Acanthoscelides obtectus). Evol Ecol. 2004 Mar 1;18(2):187.

58. Arnqvist G, Stojković B, Rönn JL, Immonen E. The pace-of-life: A sex-specific link between metabolic rate and life history in bean beetles. Funct Ecol [Internet]. 2017 [cited 2019 Mar 6];31(12):2299–309. Available from: https://besjournals.onlinelibrary.wiley.com/doi/abs/10.1111/1365-2435.12927

59. van de Pol M, Verhulst S. Age-Dependent Traits: A New Statistical Model to Separate Within- and Between-Individual Effects. Am Nat. 2006 May 1;167(5):766–73.

60. Grieshop K, Stångberg J, Martinossi-Allibert I, Arnqvist G, Berger D. Strong sexual selection in males against a mutation load that reduces offspring production in seed beetles. J Evol Biol [Internet]. 2016 Jun [cited 2019 Jan 17];29(6):1201–10. Available from: http://doi.wiley.com/10.1111/jeb.12862

61. Biemont JC. Effects of the irradiations x on the reproductive power of Acanthoscelides obtectus Say (coleoptera: bruchidae). Ann Zool Ecol Anim V 5 No 4 Pp 581–591 [Internet]. 1973 Jan 1 [cited 2022 Feb 24]; Available from: https://www.osti.gov/biblio/4119273-effects-irradiations-reproductive-power-acanthoscelides-obtectus-say-coleoptera-bruchidae

62. Kempner ES. Effects of high-energy electrons and gamma rays directly on protein molecules. J Pharm Sci. 2001 Oct;90(10):1637–46.

63. Dowling DK, Simmons LW. Reactive oxygen species as universal constraints in life-history evolution. Proc R Soc B Biol Sci [Internet]. 2009 May 22 [cited 2019 Jan 17];276(1663):1737–45. Available from: http://www.royalsocietypublishing.org/doi/10.1098/rspb.2008.1791

64. Shabalina SA, Yampolsky LY, Kondrashov AS. Rapid decline of fitness in panmictic populations of Drosophila melanogaster maintained under relaxed natural selection. Proc Natl Acad Sci [Internet]. 1997 Nov 25 [cited 2018 Feb 19];94(24):13034–9. Available from: http://www.pnas.org/content/94/24/13034

65. Hadfield JD. MCMC Methods for Multi-Response Generalized Linear Mixed Models: The MCMCglmm R Package. J Stat Softw. 2010;33(2):1–22.

66. R Core Team. R: A Language and Environment for Statistical Computing [Internet]. Vienna, Austria: R Foundation for Statistical Computing; 2020. Available from: https://www.R-project.org/

67. Wickham H. ggplot2: Elegant Graphics for Data Analysis [Internet]. Springer-Verlag New York; 2016. Available from: https://ggplot2.tidyverse.org

68. Maklakov AA, Immler S, Lovlie H, Flis I, Friberg U. The effect of sexual harassment on lethal mutation rate in female Drosophila melanogaster. Proc R Soc B Biol Sci [Internet]. 2012 Nov 21 [cited 2019 Jan 17];280(1750):20121874–20121874. Available from: http://rspb.royalsocietypublishing.org/cgi/doi/10.1098/rspb.2012.1874

69. Bartosch -Harlid A, Berlin S, Smith NGC, Mosller AP, Ellegren H. Life History and the Male Mutation Bias. Evolution [Internet]. 2003 Oct 1 [cited 2018 Dec 11];57(10):2398–406. Available from: https://onlinelibrary.wiley.com/doi/abs/10.1111/j.0014-3820.2003.tb00251.x

70. Bromham L. The genome as a life-history character: why rate of molecular evolution varies between mammal species. Philos Trans R Soc B Biol Sci [Internet]. 2011 Sep 12 [cited 2019 Jan 17];366(1577):2503–13. Available from: http://rstb.royalsocietypublishing.org/cgi/doi/10.1098/rstb.2011.0014

71. Ellegren H. Characteristics, causes and evolutionary consequences of male-biased mutation. Proc R Soc B Biol Sci [Internet]. 2007 Jan 7 [cited 2022 Feb 6];274(1606):1–10. Available from: https://royalsocietypublishing.org/doi/10.1098/rspb.2006.3720

72. Stojković B, Đorđević M, Janković J, Savković U, Tucić N. Heterosis in age-specific selected populations of a seed beetle: sex differences in longevity and reproductive behavior. Insect Sci. 2015 Apr;22(2):295–309.

73. Nussey DH, Kruuk LEB, Donald A, Fowlie M, Clutton-Brock TH. The rate of senescence in maternal performance increases with early-life fecundity in red deer. Ecol Lett. 2006;9(12):1342–50.

74. Jones OR, Scheuerlein A, Salguero-Gómez R, Camarda CG, Schaible R, Casper BB, et al. Diversity of ageing across the tree of life. Nature [Internet]. 2014 Jan [cited 2023 Oct 8];505(7482):169–73. Available from: https://www.nature.com/articles/nature12789

75. Schärer L, Rowe L, Arnqvist G. Anisogamy, chance and the evolution of sex roles. Trends Ecol Evol [Internet]. 2012 May [cited 2019 Jan 17];27(5):260–4. Available from: https://linkinghub.elsevier.com/retrieve/pii/S0169534712000043

76. Servedio MR, Powers JM, Lande R, Price TD. Evolution of sexual cooperation from sexual conflict. Proc Natl Acad Sci [Internet]. 2019 Nov 12 [cited 2020 Feb 13];116(46):23225–31. Available from: https://www.pnas.org/content/116/46/23225

77. Hurst LD, Ellegren H. Sex biases in the mutation rate. Trends Genet. 1998 Nov 1;14(11):446–52.

78. Sayres MAW, Makova KD. Genome analyses substantiate male mutation bias in many species. BioEssays [Internet]. 2011 [cited 2019 Jan 18];33(12):938–45. Available from: https://onlinelibrary.wiley.com/doi/abs/10.1002/bies.201100091

79. Gao Z, Moorjani P, Sasani TA, Pedersen BS, Quinlan AR, Jorde LB, et al. Overlooked roles of DNA damage and maternal age in generating human germline mutations. Proc Natl Acad Sci. 2019 May 7;116(19):9491–500.

80. Wang RJ, Thomas GWC, Raveendran M, Harris RA, Doddapaneni H, Muzny DM, et al. Paternal age in rhesus macaques is positively associated with germline mutation accumulation but not with measures of offspring sociability. Genome Res. 2020 Jan 6;30(6):826–34.

81. Beck CW, Promislow DEL. Evolution of Female Preference for Younger Males. PLoS ONE [Internet]. 2007 Sep 26 [cited 2019 Jun 3];2(9). Available from: https://www.ncbi.nlm.nih.gov/pmc/articles/PMC1976549/

82. Segami JC, Lind MI, Qvarnström A. Should females prefer old males? Evol Lett [Internet]. 2021 [cited 2022 Feb 6];5(5):507–20. Available from: https://onlinelibrary.wiley.com/doi/abs/10.1002/evl3.250

83. Jong G de, Noordwijk AJ van. Acquisition and Allocation of Resources: Genetic (CO) Variances, Selection, and Life Histories. Am Nat [Internet]. 1992;139(4):749–70. Available from: http://www.jstor.org/stable/2462620

84. Immler S. Haploid Selection in “Diploid” Organisms. Annu Rev Ecol Evol Syst [Internet]. 2019 [cited 2021 Jun 21];50(1):219–36. Available from: 10.1146/annurev-ecolsys-110218-024709

85. 85. Sutter A, Immler S. Within-ejaculate sperm competition. Philos Trans R Soc B Biol Sci [Internet]. 2020 Dec 7 [cited 2021 Jun 21];375(1813):20200066. Available from: https://royalsocietypublishing.org/doi/full/10.1098/rstb.2020.0066

86. Rönn JL, Katvala M, Arnqvist G. Interspecific variation in ejaculate allocation and associated effects on female fitness in seed beetles. J Evol Biol [Internet]. 2008 [cited 2022 Sep 15];21(2):461–70. Available from: https://onlinelibrary.wiley.com/doi/abs/10.1111/j.1420-9101.2007.01493.x

87. Khandwala YS, Zhang CA, Lu Y, Eisenberg ML. The age of fathers in the USA is rising: an analysis of 168 867 480 births from 1972 to 2015. Hum Reprod. 2017 Oct 1;32(10):2110–6.

88. Mills M, Rindfuss RR, McDonald P, te Velde E, on behalf of the ESHRE Reproduction and Society Task Force. Why do people postpone parenthood? Reasons and social policy incentives. Hum Reprod Update. 2011 Nov 1;17(6):848–60.

