## Supplementary Tables 1 and 2 for "Coevolution of longevity and female germline maintenance"

**Supplementary Table 1a:** Number of successful and unsuccessful F0 matings

*### F0 Matings set up with Irradiated males resulting in offspring*

| | | Line | | | | | | | |
|---|---|---|---|---|---|---|---|---|---|
| | date | 1 | 2 | 3 | 4 | 5 | 6 | 7 | 8 |
| IRR 1 | 13 | 11 | 7 | 9 | 2 | 14 | 14 | 14 | 20 |
| | 14 | 10 | 12 | 11 | 7 | 13 | 22 | 16 | 17 |
| IRR 2 | 15 | 10 | 14 | 11 | 9 | 14 | 15 | 13 | 13 |
| | 16 | 16 | 11 | 12 | 14 | 19 | 15 | 19 | 23 |
| IRR3 | 17 | 18 | 16 | 14 | 13 | 21 | 20 | 20 | 18 |
| | 18 | 22 | 24 | 13 | 14 | 24 | 20 | 27 | 18 |

*### F0 Matings set up with Control males resulting in offspring*

| | | Line | | | | | | | |
|---|---|---|---|---|---|---|---|---|---|
| | date | 1 | 2 | 3 | 4 | 5 | 6 | 7 | 8 |
| IRR 1 | 13 | 6 | 5 | 7 | 4 | 7 | 10 | 10 | 10 |
| | 14 | 7 | 8 | 8 | 8 | 10 | 9 | 6 | 13 |
| IRR 2 | 15 | 8 | 7 | 5 | 4 | 3 | 14 | 7 | 2 |
| | 16 | 6 | 12 | 14 | 4 | 11 | 14 | 13 | 13 |
| IRR 3 | 17 | 9 | 10 | 8 | 12 | 9 | 17 | 16 | 15 |
| | 18 | 13 | 9 | 9 | 10 | 14 | 10 | 13 | 18 |

*### F0 Matings set up with Irradiated males resulting in no offspring*

| | | Line | | | | | | | |
|---|---|---|---|---|---|---|---|---|---|
| | date | 1 | 2 | 3 | 4 | 5 | 6 | 7 | 8 |
| IRR 1 | 13 | 13 | 5 | 8 | 9 | 6 | 7 | 9 | 6 |
| | 14 | 8 | 6 | 13 | 5 | 10 | 4 | 9 | 9 |
| IRR 2 | 15 | 15 | 24 | 17 | 14 | 14 | 9 | 10 | 15 |
| | 16 | 7 | 14 | 11 | 12 | 15 | 12 | 5 | 11 |
| IRR 3 | 17 | 11 | 12 | 14 | 14 | 22 | 19 | 23 | 26 |
| | 18 | 9 | 6 | 13 | 13 | 20 | 17 | 19 | 21 |

*### F0 Matings set up with Irradiated males resulting in no offspring*

| | | Line | | | | | | | |
|---|---|---|---|---|---|---|---|---|---|
| | date | 1 | 2 | 3 | 4 | 5 | 6 | 7 | 8 |
| IRR 1 | 13 | 4 | 5 | 3 | 3 | 5 | 5 | 3 | 3 |
| | 14 | 6 | 3 | 4 | 2 | 6 | 8 | 6 | 5 |
| IRR 2 | 15 | 7 | 6 | 10 | 9 | 11 | 5 | 9 | 11 |
| | 16 | 10 | 3 | 4 | 9 | 6 | 6 | 9 | 10 |
| IRR 3 | 17 | 7 | 8 | 11 | 6 | 17 | 11 | 12 | 11 |
| | 18 | 9 | 8 | 9 | 3 | 13 | 6 | 11 | 11 |

*Base males were irradiated (or kept as controls) on three occasions (13$^{th}$, a5th and 17$^{th}$) and mated to females on the same day or the following day.*

*Line 1-4: E regime*

*Line 5-8: L regime.*

**Supplementary Table 1b:** Samples sizes per treatment in analyses

*# Number of F0 females mated to control males and scored for reproductive output as young or aged.*

| | Young | Aged |
|---|---|---|
| E1 | 21 | 12 |
| E2 | 22 | 13 |
| E3 | 19 | 8 |
| E4 | 20 | 9 |
| L5 | 14 | 18 |
| L6 | 20 | 21 |
| L7 | 17 | 15 |
| L8 | 19 | 23 |

*#Number of F1 couples scored for F2 offspring production to measure germline maintenance*

Control treatment

| | aged | young |
|---|---|---|
| E1 | 6 | 14 |
| E2 | 8 | 15 |
| E3 | 6 | 12 |
| E4 | 6 | 12 |
| L5 | 13 | 5 |
| L6 | 12 | 12 |
| L7 | 11 | 10 |
| L8 | 13 | 8 |

Irradiated treatment

| | aged | young |
|---|---|---|
| E1 | 16 | 25 |
| E2 | 17 | 27 |
| E3 | 14 | 23 |
| E4 | 11 | 23 |
| L5 | 19 | 19 |
| L6 | 20 | 14 |
| L7 | 27 | 24 |
| L8 | 25 | 23 |

**Supplementary table 2a:** Reproductive schedule of F0 females mated to ctrl males

DIC: 2040.019

| | post.mean | l-95% CI | u-95% CI | eff.samp |
|---|---|---|---|---|
| line | 0.005718 | 1.364e-07 | 0.02989 | 2000 |

| | post.mean | l-95% CI | u-95% CI | eff.samp |
|---|---|---|---|---|
| line:age | 0.004063 | 1.498e-07 | 0.02253 | 1994 |

| | post.mean | l-95% CI | u-95% CI | eff.samp |
|---|---|---|---|---|
| date | 0.01496 | 1.391e-07 | 0.07351 | 1676 |

| | post.mean | l-95% CI | u-95% CI | eff.samp |
|---|---|---|---|---|
| regimeL.units | 1.0024 | 0.7667 | 1.2797 | 2000 |
| regimeE.units | 0.7315 | 0.4845 | 0.9626 | 2000 |

Location effects: offspring ~ regime * class + male1

| | post.mean | l-95% CI | u-95% CI | eff.samp | pMCMC |
|---|---|---|---|---|---|
| Intercept(Young L) | 2.6795 | 2.4157 | 2.9847 | 2000 | <5e-04 *** |
| Regime.E | 0.2111 | -0.1164 | 0.4998 | 2000 | 0.188 |
| Aged | 0.1372 | -0.1590 | 0.4555 | 2000 | 0.396 |
| maleAge | 0.2032 | -0.0097 | 0.4233 | 2000 | 0.076 . |
| **regime.E:Aged** | **-0.8458** | **-1.3391** | **-0.3858** | **2000** | **<5e-04 *** ** |

**Supplementary table 2b:** Germline maintenance of F0 females based on analysis of F2 adult offspring production

DIC: 3448.681

| | post.mean | l-95% CI | u-95% CI | eff.samp |
|---|---|---|---|---|
| line | 0.00141 | 1.211e-07 | 0.007541 | 2000 |

| | post.mean | l-95% CI | u-95% CI | eff.samp |
|---|---|---|---|---|
| line:age | 0.001285 | 1.187e-07 | 0.007419 | 2000 |

| | post.mean | l-95% CI | u-95% CI | eff.samp |
|---|---|---|---|---|
| line:radiation | 0.001844 | 1.181e-07 | 0.009375 | 2000 |

| | post.mean | l-95% CI | u-95% CI | eff.samp |
|---|---|---|---|---|
| line:radiation:age | 0.002043 | 1.047e-07 | 0.01102 | 2000 |

| | post.mean | l-95% CI | u-95% CI | eff.samp |
|---|---|---|---|---|
| regimeE:ctrl.units | 0.4375 | 0.2703 | 0.6271 | 2000 |
| regimeL:ctrl.units | 0.3329 | 0.2086 | 0.4709 | 2000 |
| regimeE:irradiated.units | 0.9043 | 0.6516 | 1.1610 | 2000 |
| regimeL:irradiated.units | 1.0935 | 0.8062 | 1.4061 | 2000 |

Location effects: offspring ~ regime * class * rad

| | post.mean | l-95% CI | u-95% CI | eff.samp | pMCMC | |
|---|---|---|---|---|---|---|
| Intercept(aged E Ctrl) | 3.46663 | 3.18391 | 3.74447 | 2087 | <5e-04 | *** |
| RegimeL | -0.04728 | -0.39223 | 0.28763 | 2003 | 0.791 | |
| young | -0.05963 | -0.36988 | 0.30204 | 2000 | 0.744 | |
| **irradiated** | **-0.54281** | **-0.94203** | **-0.15079** | **2020** | **0.004** | ** |
| regimeL:young | 0.37427 | -0.04951 | 0.79023 | 2000 | 0.082 | . |
| regimeL:irradiated | 0.28412 | -0.19595 | 0.79187 | 1800 | 0.247 | |
| young:irradiated | 0.19963 | -0.32662 | 0.67202 | 2000 | 0.401 | |
| **regimeL:young:irradiated** | **-0.87734** | **-1.51822** | **-0.23396** | **2000** | **0.008** | ** |
